# Utilizing a PTPN22 gene signature to predict response to targeted therapies in rheumatoid arthritis

**DOI:** 10.1101/586982

**Authors:** Hui-Hsin Chang, Ching-Huang Ho, Beverly Tomita, Andrea A. Silva, Jeffrey A. Sparks, Elizabeth W. Karlson, Deepak A. Rao, Yvonne C. Lee, I-Cheng Ho

## Abstract

Despite the development of several targeted therapies for rheumatoid arthritis (RA), there is still no reliable drug-specific predictor to assist rheumatologists in selecting the most effective targeted therapy for each patient. Recently, a gene signature caused by impaired induction of PTPN22 in anti-CD3 stimulated peripheral blood mononuclear cells (PBMC) was observed in healthy at-risk individuals. However, the downstream target genes of PTPN22 and the molecular mechanisms regulating its expression are still poorly understood. Here we report that the PTPN22 gene signature is also present in PBMC from patients with active RA and can be reversed after effective treatment. The expression of PTPN22 correlates with that of more than 1000 genes in Th cells of anti-CD3 stimulated PBMC of healthy donors and is inhibited by TNFα or CD28 signals, but not IL-6, through distinct mechanisms. In addition, the impaired induction of PTPN22 in PBMC of patients with active RA can be normalized in vitro by several targeted therapies. More importantly, the in vitro normalization of PTPN22 expression correlates with clinical response to the targeted therapies in a longitudinal RA cohort. Thus, in vitro normalization of PTPN22 expression by targeted therapies can potentially be used to predict clinical response in a drug-specific manner.

## 1. Introduction

Rheumatoid arthritis (RA), a disease featuring joint inflammation and the presence of anti-citrullinated protein antibodies (ACPA) [1], affects 1-2% of the general population in North America. Despite the development of several targeted therapies, there is no reliable predictor to assist rheumatologists in choosing the most effective targeted therapy for each individual patient. Therefore, finding molecular-based predictors of differential response to targeted therapies remains a research priority of the American College of Rheumatology and European League Against Rheumatism (https://www.rheumatology.org/Portals/0/Files/ACR-Research-Agenda-2016-2020.pdf) [2].

The underlying causes of RA are still not fully understood. Genome-wide association studies have identified more than 100 genetic loci that are associated with higher risk of RA [3]. A C-to-T single nucleotide polymorphism (C1858T SNP) located at the position 1858 of human PTPN22 cDNA carries the highest risk of RA among all non-HLA genetic variations. PTPN22 is a non-receptor protein tyrosine phosphatase expressed preferentially in hematopoietic cells. Its phosphatase activity has been shown to attenuate the activation signals in lymphocytes [4, 5], modulate the polarization of macrophages [6], and suppress the activation of NLRP3 inflammasome in myeloid cells [7]. Interestingly, PTPN22 also augments TLR-induced production of type 1 interferon but suppresses protein citrullination independently of its phosphatase activity [8, 9], suggesting the presence of phosphatase-independent functions.

Another major risk factor of RA is family history [10]. Recently, we discovered that PBMC from a great majority of healthy first-degree relatives (FDRs) of RA patients displayed a molecular signature characterized by impaired induction of PTPN22 by anti-CD3, attenuated expression of PFKFB3 and ATM, but heightened expression of Th17 cytokines [11]. We further demonstrated that the impaired induction of PTPN22 was at the root of the signature, henceforth referred to as the PTPN22 gene signature. Interestingly, this signature predates the onset of ACPA and is independent of the C1858T SNP of PTPN22. However, the downstream target genes of PTPN22 and the molecular mechanism regulating its expression are still poorly understood.

In this work, we determine the upstream regulators of PTPN22 and identify more than 1000 genes, whose expression correlates with that of PTPN22. We further demonstrate that the PTPN22 gene signature is also present in PBMC from patients with active RA and is reversed after effective treatments. In addition, the PTPN22 gene signature can be reversed in vitro by etanercept, adalimumab, or tocilizumab. More importantly, the in vitro reversal of the PTPN22 gene signature correlates with clinical response to the targeted therapies.

## 2. Material and methods

### 2.1 Human subjects

PBMCs were obtained from the following sources:

1. Partners HealthCare Biobank: an enterprise biobank of consented patients’ samples at Partners HealthCare consented for research studies using genetics and potential for re-contact for specific studies (Massachusetts General Hospital and Brigham and Women’s Hospital, Boston, MA).
2. Central Pain in Rheumatoid Arthritis (CPIRA) study [12]: a multicenter, prospective, observational study designed to examine the relationship between pain and treatment response in RA. At enrollment, subjects had active RA and were initiating or switching disease modifying antirheumatic drugs.
3. Profiling of Cell Subsets in Human Diseases (PROSET-HD): a research initiative of Brigham and Women’s Hospital comparing immune cells in the blood from patients with or without inflammatory diseases.

### 2.2 Study approval

This study has been approved by Partners Human Research Committee (PHRC), Boston, MA. Informed consent was obtained from participants prior to inclusion to the studies.

### 2.3 Purification and stimulation of PBMCs

PBMCs were isolated from whole human peripheral blood by Ficoll-Paque PLUS (17-1440-03, GE Healthcare, Pittsburgh, PA) density gradient centrifugation and cryopreserved prior to use. PBMCs were then thawed and plated in 24-well plates (2-2.5 millions/1ml/well) pre-coated with anti-CD3 (2.5 ug/ml, HIT3a clone, Biolegend, San Diego, CA) for 24 hours before harvesting. In some experiments, abatacept, etanercept, adalimumab, anti-CD28 (Cat. #302914, Biolegend), or TNFα (PHC3015, Life Technology, Carlsbad, CA) was added during the stimulation.

### 2.4 Purification of Th and non-Th cells

Th (CD4+) cells and non-Th (CD4-) cells were sorted from PBMC using a BD FACSAria cell sorter after staining with anti-human CD4 PE-conjugated antibody (Cat. #317410, Biolegend) or separated by human CD4+ T cell isolation kit (Cat. #130-045-101, Miltenyi).

### 2.5 RNA-seq and data analysis

1000 cells of CD4+ cells were sorted directly into 5 ul of TCL buffer (Qiagene, Cat#1031576) in skirted Eppendorf 96-well twin.tec PCR plate (Eppendorf). Smart-seq2 modified from a published method [13] was carried out at Broad Technology Labs. RNA-seq data was analyzed with Qlucore Omics Explorer (https://qlucore.com) to identify differentially expressed genes, which were further analyzed with Ingenuity Pathway Analysis (https://www.qiagenbioinformatics.com/products/ingenuity-pathway-analysis/) to identify over-representative pathways and upstream regulators.

### 2.6 Quantitative RNA analysis

RNA isolation, reverse transcription, and quantitative PCR (qPCR) were performed as described [14]. The transcript level thus detected was normalized against that of actin from the same sample. Unless indicated otherwise, the average of the normalized values from the control group in each figure was arbitrarily set as 1. The sequences of the primers used in qPCR are listed in Supplemental Table 1.

### 2.7 Statistical analyses

Statistical analyses were performed with two-tailed Student’s t test (Figure 1B, 1C, 2, and Supplemental Figure 4A), ratio paired two-tailed Student’s t test (Figure 1D, 3, 5,and 6), and Pearson correlation test (Figure 4B, 4C, and Supplemental Figure 4B). A p-value < 0.05 was considered significant

**Figure 1.**
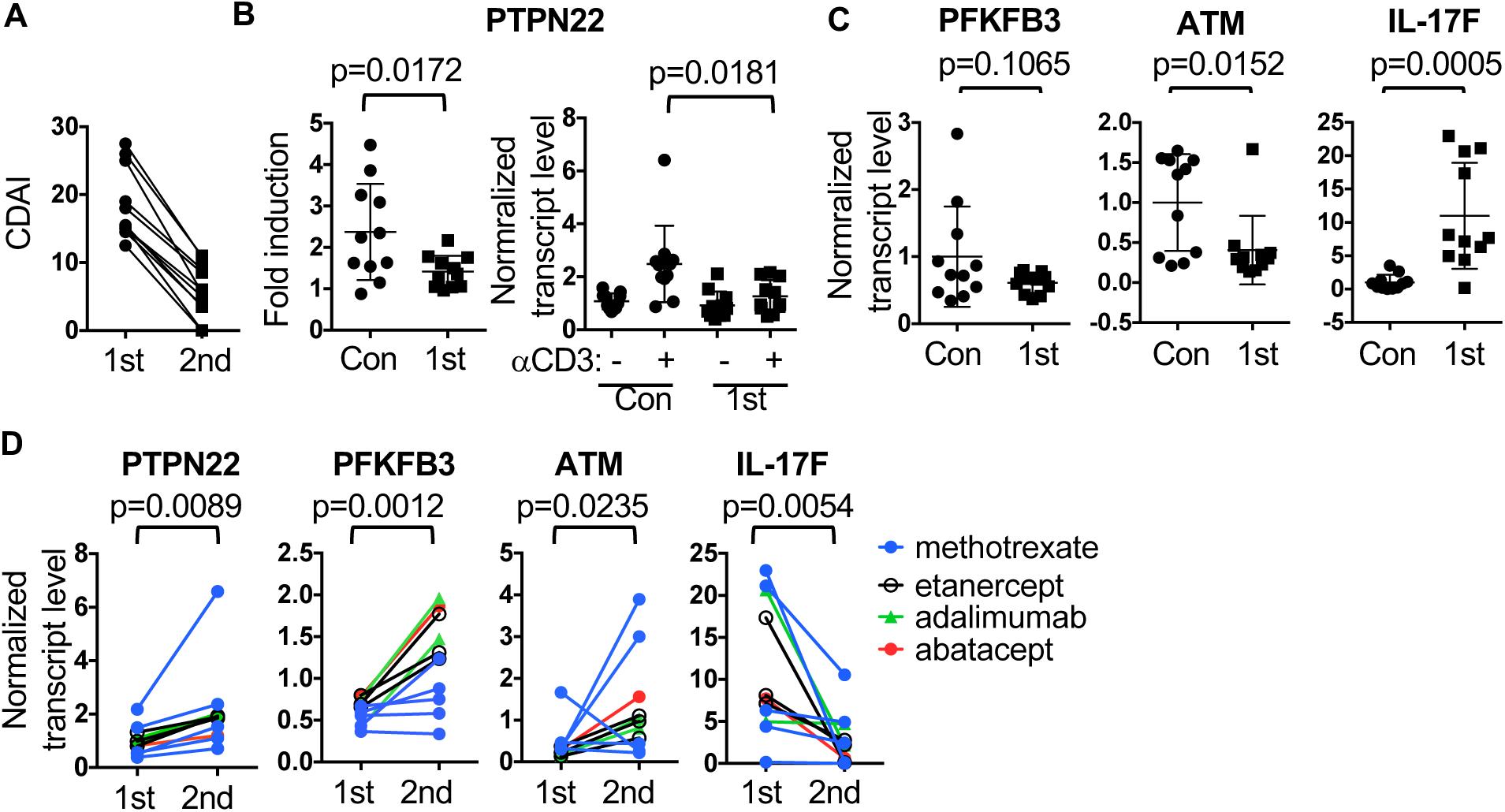
Detection of the PTPN22 gene signature in active RA patients. The 1st and 2nd visit CDAI scores of 11 RA patients are shown in **A**. Their paired PBMC (1st visit and 2nd visit) and PBMC from 11 healthy donors were stimulated with anti-CD3. The anti-CD3-induced fold change in PTPN22 (**B**, the left panel) and the normalized levels of PTPN22 (**B**, the right panel), PFKFB3 (**C**), ATM (**C**), and IL-17F (**C**) from control PBMC and 1st visit PBMC of RA subjects are shown. The normalized levels of PTPN22, PFKFB3, ATM, and IL-17F from anti-CD3 stimulated 1st visit and 2nd visit PBMC of RA subjects are shown in **D**. The data points from the same RA subjects are connected with lines. The healthy donors were used as the control group for normalization of qPCR data in **B**-**D**.

**Figure 2.**
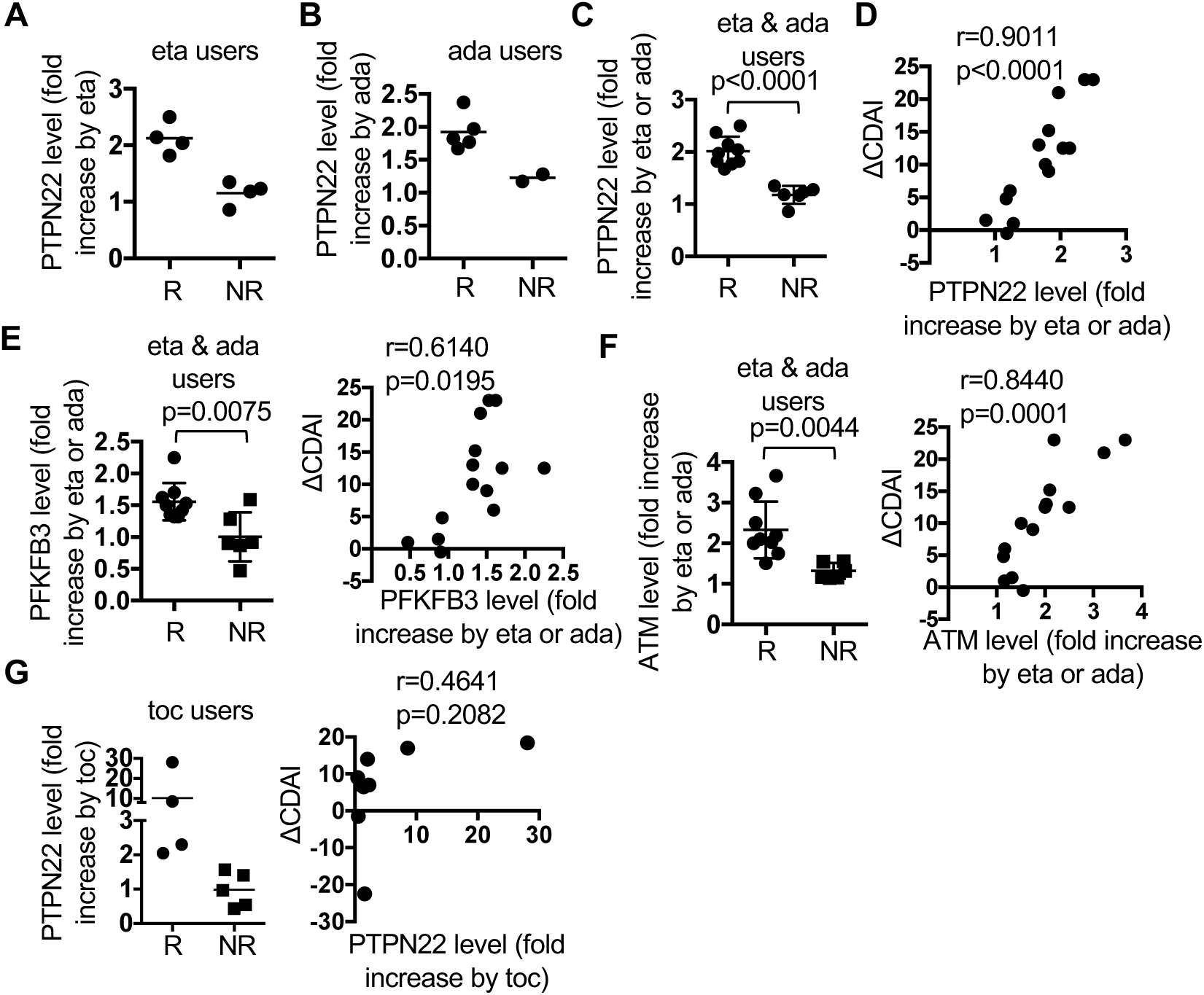
In vitro reversal of the PTPN22 gene signature by TNFα inhibitors or tocilizumab correlates with clinical response. PBMC from 23 active RA subjects were stimulated with anti-CD3 in the presence of etanercept (eta, for etanercept users, **A** & **C-F**), adalimumab (ada, for adalimumab users, **B-F**), or tocilizumab (toc, for tocilizumab users, **G**). The fold increase in the level of PTPN22 by etanercept, adalimumab, or tocilizumab in responders and non-responders is shown in **A**, **B**, and the left panel of **G**. The data from **A** and **B** are combined into **C** and plotted against ΔCDAI in **D**. The fold increase in the transcript level of PFKFB3 (**E**) and ATM (**F**) by etanercept or adalimumab is shown in the left panels of **E** and **F**, and plotted against ΔCDAI (the right panels of **E** and **F**). The fold increase in the transcript level of PTPN22 by tocilizumab is also plotted against ΔCDAI (the right panel of **G**).

**Figure 3.**
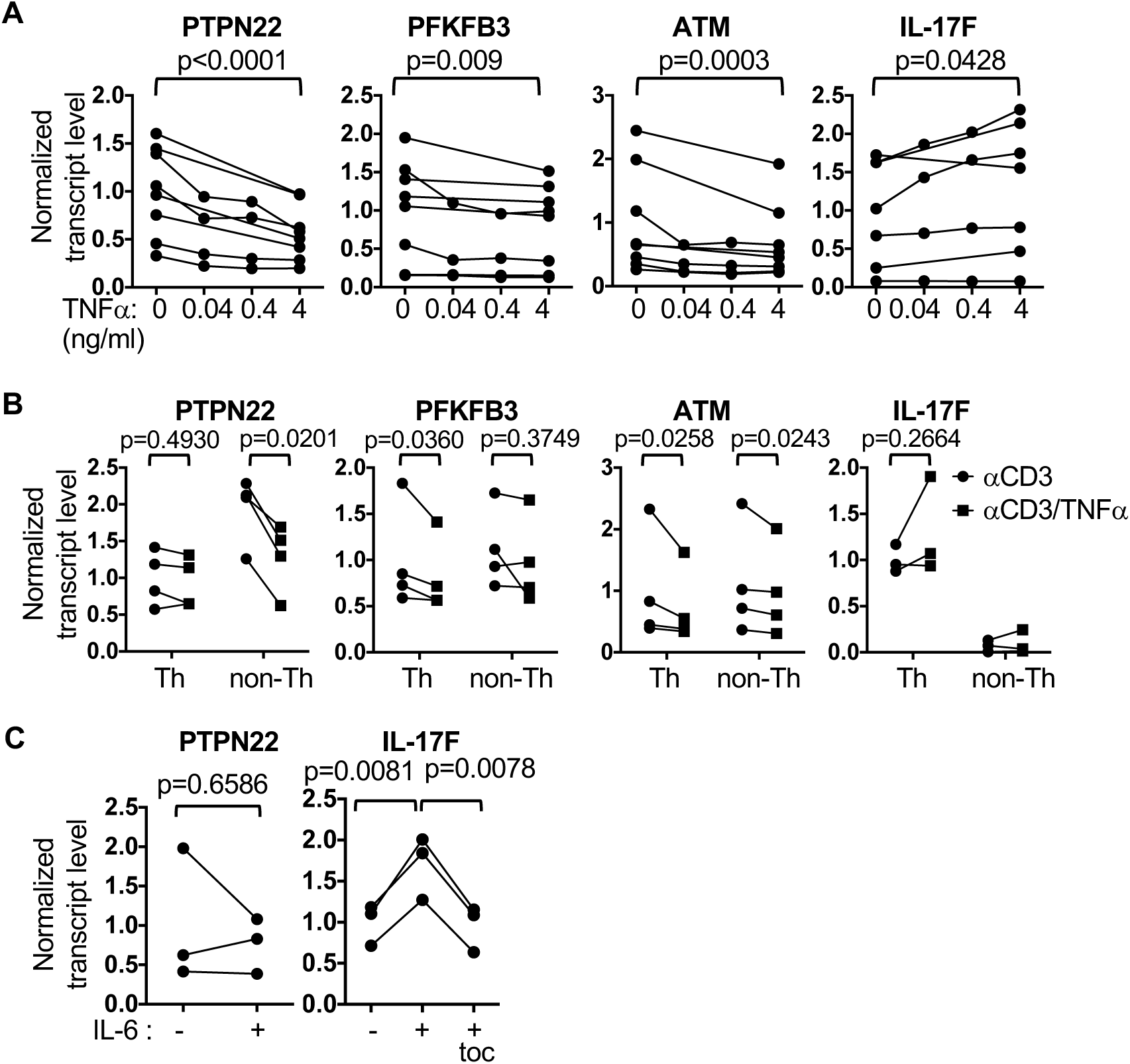
Recapitulation of the PTPN22 gene signature with TNFα but not IL-6. PBMC from healthy individuals were stimulated with anti-CD3 in the absence or presence of TNFα (0.04 - 4 ng/ml, **A** & **B**) or IL-6 (50 ng/ml, **C**) for 24 hours. Tocilizumab (toc) at 100 ug/ml was added along with IL-6 in some samples (**C**). A fraction of the stimulated PBMC were separated into CD4+ Th or CD4-non-Th cells (**B**). The transcript levels of indicated genes in PBMC (**A** & **C**) and indicated populations (**B**) were determined with qPCR. Paired Student t test was used to compare 0 ng/ml and 4 ng/ml of TNFα. Anti-CD3 only groups were used as the control groups for normalization of qPCR data. Data points from the same donors were connected with lines.

**Figure 4.**
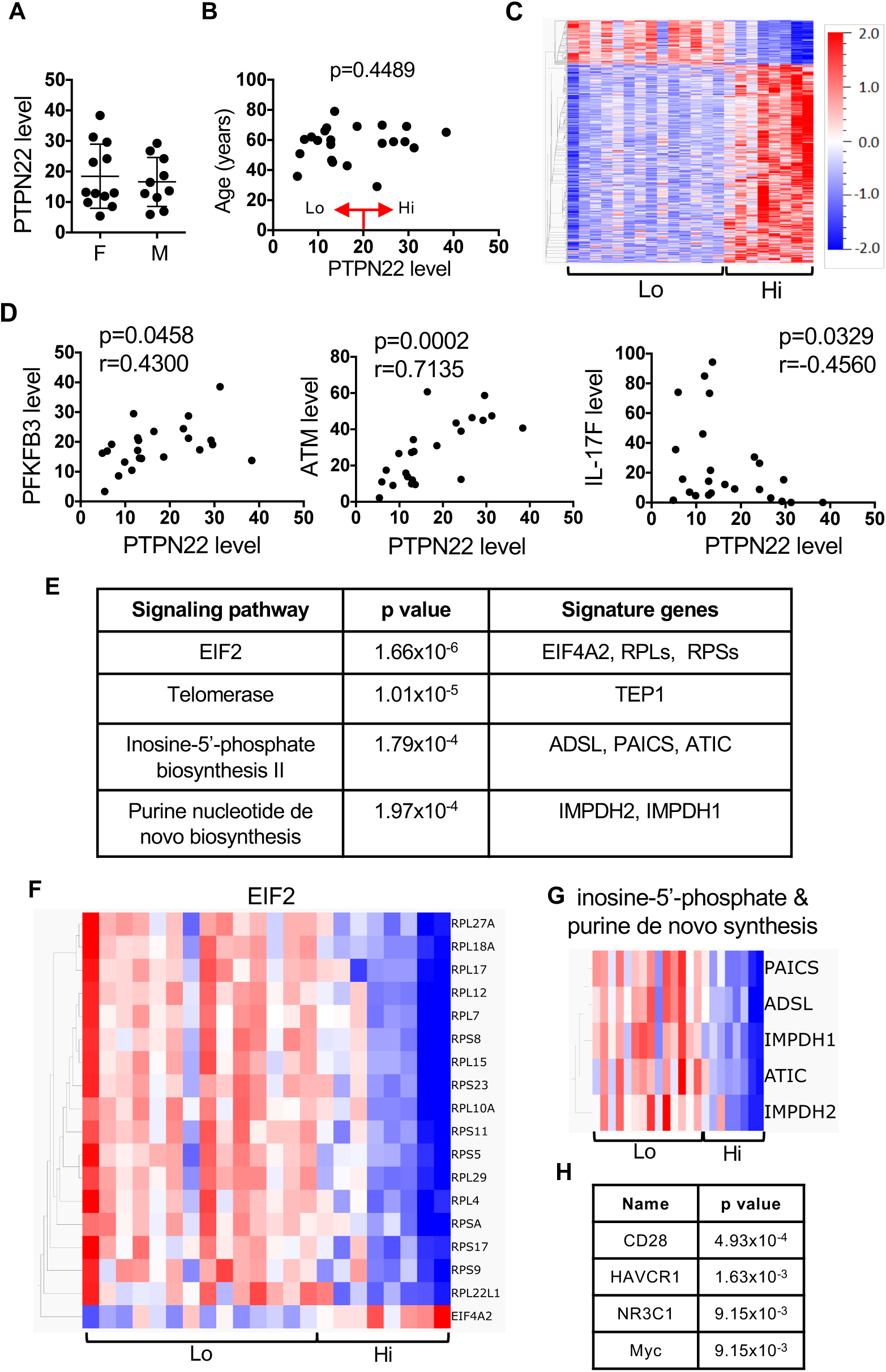
Identification of potential downstream target genes and upstream regulators of PTPN22 in Th cells. PBMC from 22 healthy individuals were stimulated with anti-CD3 for 24 hours. CD4+ T cells were sorted and subjected to RNA-seq analysis. The number of read of PTPN22 transcript was used as surrogate of its transcript level. The levels in female and male are shown in **A**, are plotted against age in **B**. Genes that are differentially expressed between PTPN22hi and PTPN22lo group were identified with Qlucore. The heat map of the differentially expressed gene is show in **C**. The correlations between PTPN22 levels and the levels PFKFB3, ATM, and IL-17F are shown in **D**. The differentially expressed genes were analyzed with INGENUITY. The top 4 over-representative pathways and their p values and signature genes are shown in **E**. The heatmap of the signature genes of EIF2 pathway is shown in **F** and of inosine-5-phohphate biosynthesis plus purine de novo synthesis is shown in **G**. The top 4 upstream regulators and their p values predicted by INGENUITY are in **H**.

**Figure 5.**
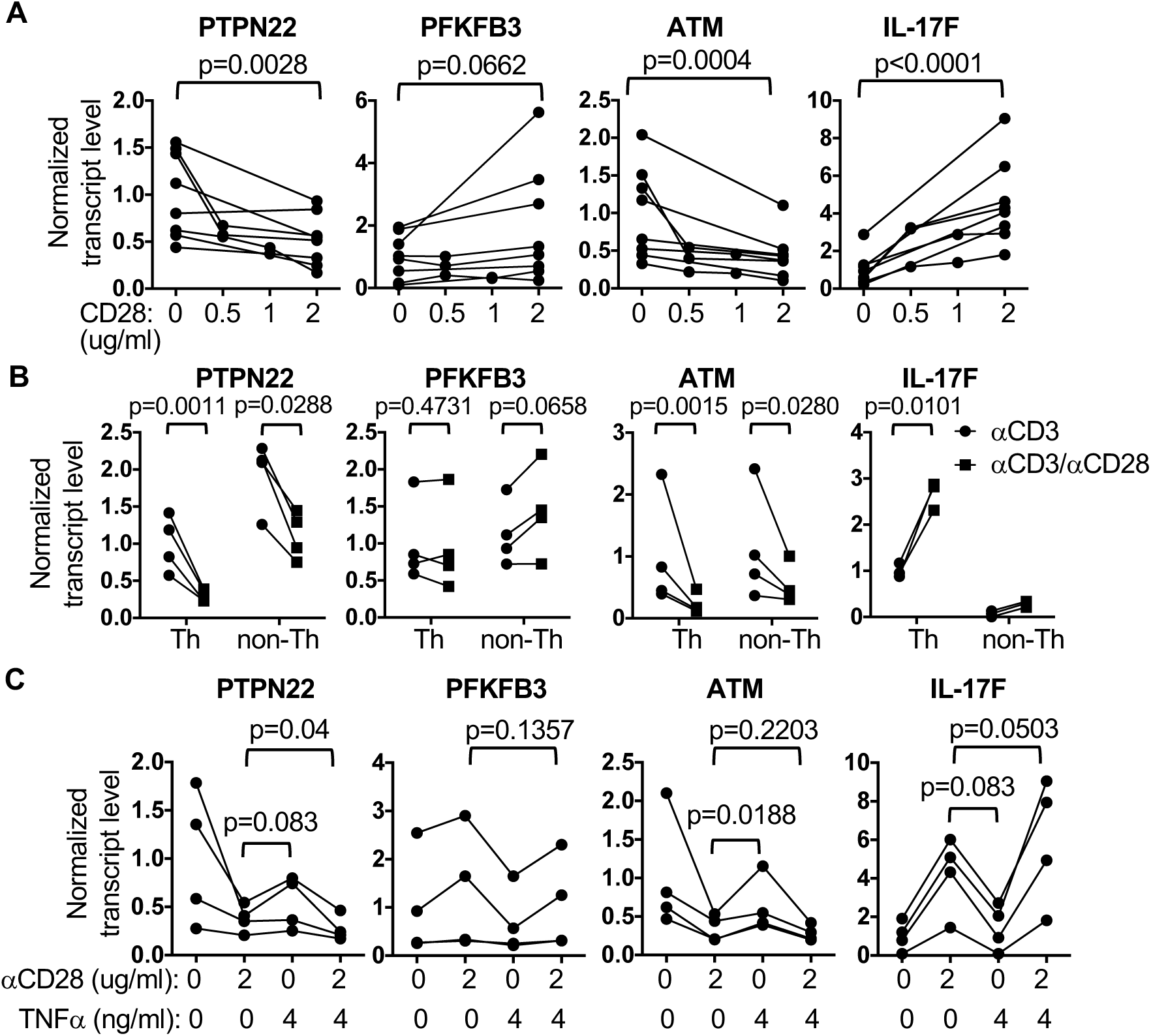
Partial recapitulation of the PTPN22 gene signature with anti-CD28. PBMC from healthy donors were stimulated with anti-CD3 for 24 hours in the absence or presence of anti-CD28 at indicated concentration (**A** & **C**) or 2 ug/ml (**B**) for 24 hours. PBMC that were stimulated in the presence of 2 ug/ml of anti-CD28 were further separated into Th and non-Th cells (**B**). TNFα was also added to some samples (**C**). Transcript levels of the indicated genes in the indicated cell populations were measured with qPCR. Statistical analysis was performed with paired Student t test. Anti-CD3 only groups were used as the control groups for normalization of qPCR data. Data points from the same donors were connected with lines.

**Figure 6.**
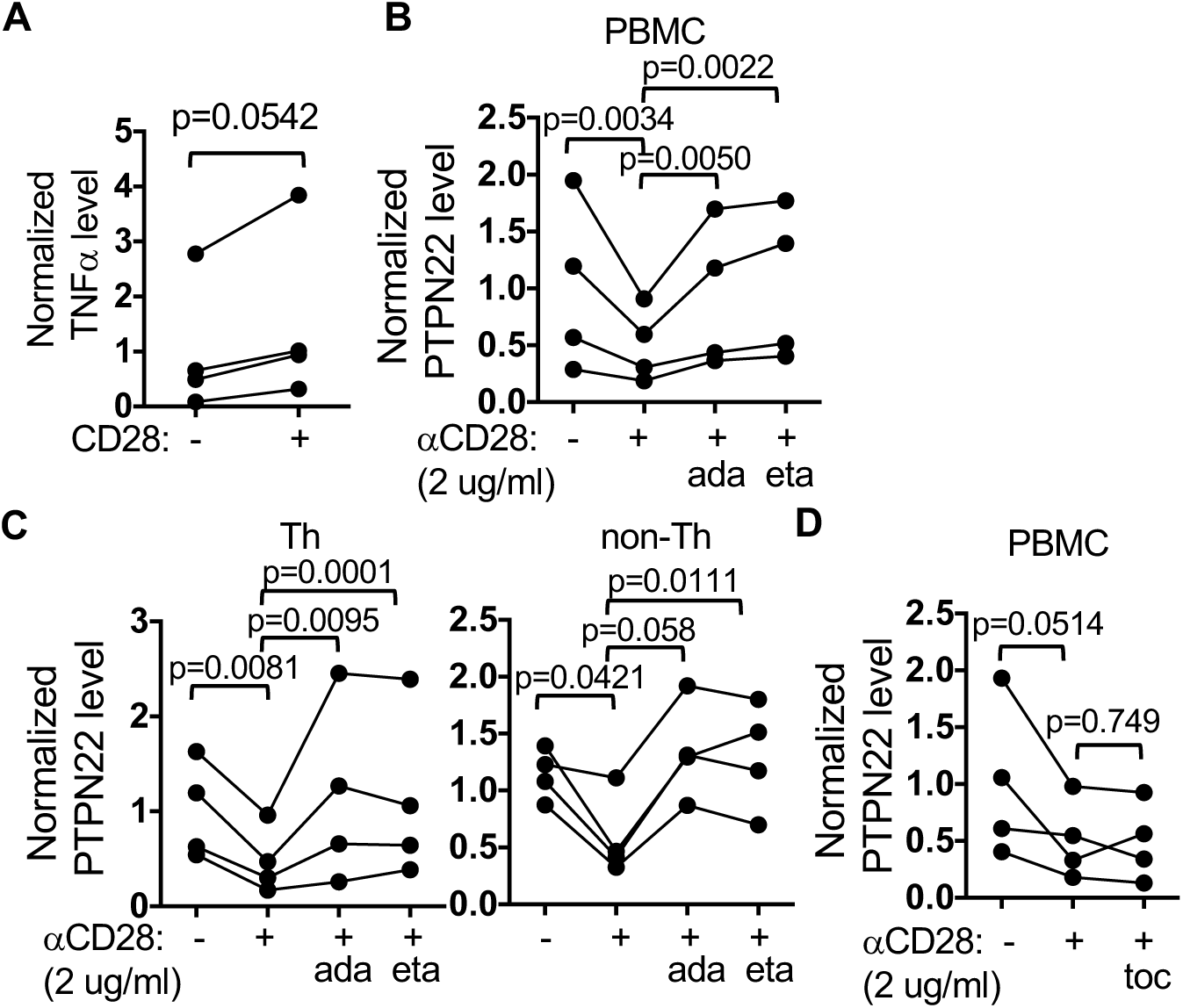
Counteracting the effect of anti-CD28 with TNFα inhibitors. PBMC from healthy donors were stimulated with anti-CD3 for 24 hours in the absence or presence of anti-CD28 (2 ug/ml) for 24 hours (**A**-**D**). Etanercept (eta, 100 ug/ml, **B** & **C**), adalimumab (ada, 100 ug/ml, **B** & **C**), or tocilizumab (toc, 100 ug/ml, **D**) was added to some samples. The samples in **B** were further separated into Th and non-Th cells (**C**). Transcript levels of the indicated genes in the indicated cell populations were measured with qPCR. Statistical analysis was performed with paired Student t test. Anti-CD3 only groups were used as the control groups for normalization of qPCR data. Data points from the same donors were connected with lines.

## 3. Results

### 3.1 Detection of the PTPN22 gene signature in active RA patients

While we have demonstrated that the PTPN22 gene signature is present in healthy at-risk individuals, it is still unclear if the gene signature is also present in PBMC of patients with active RA. If it is, can PTPN22 gene signature be normalized with effective treatment? To address these questions, we took advantage of the study cohort of CPIRA (Central Pain in Rheumatoid Arthritis). CPIRA recruited RA patients who were starting or switching to a new treatment. PBMC were collected at the time of start or switch (1st visit) and then approximately 12 weeks later (2nd visit). We selected 11 RA patients from the CPIRA cohorts who had a Clinical Disease Activity Index (CDAI) >10 in the 1st visit and had a reduction of CDAI>10 in the 2nd visit (Figure 1A). Five of the 11 RA patients started or switched to methotrexate, 3 to etanercept, 2 to adalimumab, and 1 to abatacept. We also included PBMC from 11 healthy donors who did not carry the C1858T SNPs and had no family history of RA. PBMCs were stimulated with anti-CD3. In agreement with our published data [11], the expression of PTPN22 increased approximately twofold in control PBMC but not in 1st visit PBMCs of subjects with active RA (Figure 1B, the left panel). While the level of PTPN22 in resting PBMC from 1st visit of the subjects with active RA was not lower than that from control resting PBMC, the impaired induction resulted in a lower level of PTPN22 after anti-CD3 stimulation (Figure 1B, the right panel). In addition, the levels of PFKFB3 and ATM were lower, whereas the level of IL-17F was higher in subjects with active RA compared to healthy controls (Figure 1C). Furthermore, effective treatment enabled anti-CD3 to induce PTPN22 and normalized the expression of PTPN22, PFKFB2, CPIRA, and IL-17F in the RA subjects (Figure 1D). Thus, the PTPN22 gene signature is also detected in patients with active RA and can be normalized with effective treatments.

### 3.2 In vitro reversal of the PTPN22 gene signature by TNFα inhibitors

The observation that the PTPN22 gene signature is reversed after effective treatment prompted us to examine if targeted therapies, such as TNFα inhibitors (TNFis), when added in vitro are sufficient to reverse the PTPN22 gene signature in PBMC from symptomatic RA patients. In a pilot experiment, we took PBMC from two patients with active RA and stimulated the PBMC with anti-CD3 in the presence or absence of etanercept. Interestingly, we found that etanercept dose-dependently increased the level of PTPN22, PFKFB3, and ATM but decreased the level of IL-17F (Supplemental Figure 1).

### 3.3 In vitro reversal of the PTPN22 gene signature by etanercept and adalimumab correlates with clinical response

We subsequently asked if in vitro reversal of the PTPN22 gene signature by TNFis correlates with clinical response. We examined RA subjects from the CPIRA cohort who had an entry CDAI of ≥10 and started or switched to etanercept or adalimumab, the two most commonly used TNFis. Responsiveness was defined based on a published criteria [15]: a decrease of CDAI≥12 for those with baseline CDAI>22 and a decrease of CDAI≥6 for those with baseline CDAI 10-22. There were 7 etanercept users (4 responders and 3 non-responders) and 7 adalimumab users (5 responders and 2 non-responders) who met the entry criteria (Table 1). One of the adalimumab non-responders (BWH13) also had a history of non-response to etanercept before study enrollment. Their PBMCs were tested in a blinded manner in vitro with etanercept and adalimumab separately regardless of which TNFi they started or switched to. However, only the results generated from the corresponding TNFi (etanercept on etanercept users and adalimumab on adalimumab users) were used for analysis because, with the exception of subject BWH13, no information was available on the participants’ response to the other TNFi. Subject BWH13 was counted as both an adalimumab non-responder and an etanercept non-responder and was included in both sets of analyses.

**Table 1.** Characteristics of CPIRA subjects and the results from the in vitro assay described in Figure 5

| Drug | Response | ID | age | gender | RF/<br>CCP | 1st<br>CDAI | 2nd<br>CDAI | ΔCDAI | PTPN22<br>fold<br>increase<br>by eta | PTPN22<br>fold<br>increase<br>by ada | PTPN22<br>fold<br>increase<br>by toci |
| --- | --- | --- | --- | --- | --- | --- | --- | --- | --- | --- | --- |
| eta | R | BWH1 | 76 | F | + | 19 | 6.5 | 12.5 | **2.04** | 2.2 | ND |
|  |  | BWH2 | 56 | F | + | 12.5 | 0 | 12.5 | **2.14** | 1.7 | ND |
|  |  | BWH3 | 40 | F | + | 15 | 6 | 9 | **1.82** | 2 | ND |
|  |  | BWH4 | 44 | F | + | 35 | 12 | 23 | **2.5** | 1.7 | ND |
|  | NR | BWH5 | 45 | F | + | 27 | 25.5 | 1.5 | **0.86** | 1.06 | ND |
|  |  | BWH6<sup>#</sup> | 58 | F | + | 21 | 21.5 | -0.5 | **1.18** | 1.86 | ND |
|  |  | BWH7<sup>#</sup> | 50 | F | + | 29 | 23 | 6 | **1.23** | 2.25 | ND |
| ada | R | BWH8<sup>#</sup> | 24 | F | + | 15.2 | 0 | 15.2 | 1.14 | **1.82** | ND |
|  |  | BWH9<sup>#</sup> | 52 | F | - | 19 | 9 | 10 | 1.35 | **1.77** | ND |
|  |  | BWH10 | 52 | F | + | 46 | 25 | 21 | 1.8 | **1.97** | ND |
|  |  | BWH11 | 25 | F | - | 57 | 34 | 23 | 1.59 | **2.37** | ND |
|  |  | BWH12 | 52 | F | + | 31 | 18 | 13 | 1.46 | **1.67** | ND |
|  | NR | BWH13 | 54 | F | - | 29 | 24.2 | 4.8 | **1.35** | **1.17** | ND |
|  |  | BWH14<sup>#</sup> | 79 | F | - | 14 | 13 | 1 | 1.54 | **1.28** | ND |
| toc | R | BWH45 | 66 | F | + | 15 | 8 | 7 | ND | ND | 2.3 |
|  |  | BWH97 | 80 | F | - | 25 | 8 | 17 | ND | ND | 8.57 |
|  |  | BWH115 | 21 | F | + | 29.7 | 15.7 | 14 | ND | ND | 2.05 |
|  |  | BWH155 | 77 | F | + | 18.5 | 0.1 | 18.4 | ND | ND | 28.07 |
|  | NR | BWH46 | 30 | F | + | 27 | 25.5 | 1.5 | ND | ND | 1.4 |
|  |  | BWH47 | 60 | F | - | 25 | 18 | 7 | ND | ND | 0.97 |
|  |  | BWH92 | 37 | F | + | 22.5 | 24 | -1.5 | ND | ND | 0.54 |
|  |  | BWH111 | 63 | F | + | 28 | 19 | 9 | ND | ND | 0.43 |
|  |  | BWH125 | 42 | F | + | 37.5 | 60 | -22.5 | ND | ND | 1.57 |

We found that the corresponding TNFi led to a 1.67-2.50-fold increase in the level of PTPN22 in anti-CD3-stimulated PBMC from responders, but only a 0.86-1.35-fold increase in non-responders (Figure 2A and 2B). When the data from etanercept and adalimumab users were combined together, there was clean separation between responders and non-responders (Figure 2C). More intriguingly, there was a positive linear correlation between fold increase in the level of PTPN22 and ΔCDAI (1st visit CDAI - 2nd visit CDAI) (Figure 2D). Etanercept and adalimumab had similar effects on the expression of PFKFB3 and ATM (Figure 2E and 2F) but did not consistently inhibit the expression of IL-17F in responders (Supplemental Figure 2). There was also no correlation between ΔCDAI and fold reduction in IL-17F.

The CPIRA cohort also includes 9 tocilizumab users (4 responders and 5 non-responders) who had entry CDAI>10 (Table 1). To determine if tocilizumab could also reverse the PTPN22 gene signature, we stimulated their 1^st^ visit PMBC with anti-CD3 in the absence or presence of tocilizumab (toc, a humanized monoclonal IgG1 against IL-6R). We found that tocilizumab led to 2-28-fold increase in the level of PTPN22 in responders but no more than 1.6 folds in non-responders (the left panel of Figure 2G). There was also a trend of positive correlation between fold increase in the level of PTPN22 and ΔCDAI (the right panel of Figure 2G).

### 3.4 TNFα modestly inhibited the expression of PTPN22

Both adalimumab and etanercept can enhance the expression of PTPN22 in responders (Figure 2). We therefore postulated that soluble TNFα could inhibit the expression of PTPN22. Indeed, TNFα dose-dependently but modestly reduced the levels of PTPN22, PFKFB3 and ATM, but slightly increased the expression of IL-17F in anti-CD3 stimulated PBMC from healthy donors (Figure 3A). When we separated the stimulated PBMC into Th and non-Th cells, we found that TNFα inhibited the expression of ATM comparably in Th cells and non-Th cells (Figure 3B). It also inhibited the expression of PTPN22 in non-Th cells and PFKFB3 in Th cells. Surprisingly, TNFα had little impact on the expression of PTPN22 in Th cells and PFKFB3 in non-Th cells. IL-17F was expectedly expressed mainly by Th cells but not non-Th cells. The effect of TNFα on the expression of IL-17F was already subtle in PBMC and negligible in Th cells. While tocilizumab also enhanced the expression of PTPN22 in responders (Figure 2G), exogenous IL-6 had little impact on the expression of PTPN22 even though it expectedly augmented the expression of IL-17F by anti-CD3 stimulated PBMC (Figure 3C). This effect of IL-6 on the expression of IL-17F was neutralized by tocilizumab.

### 3.5 Suppression of PTPN22 expression by CD28 signals

T cells, particularly Th cells, are the immediate responders to anti-CD3 stimulation. However, soluble TNFα only modestly inhibited the expression of PTPN22 in anti-CD3 stimulated PBMC and had no impact on the level of PTPN22 in Th cells. In addition, the impact of impaired induction of PTPN22 on the gene expression profile of Th cells is still unclear. We therefore set to identify other genes/signals regulating the expression of PTPN22 and candidates of its downstream target genes in Th cells. We stimulated PBMC obtained from 22 healthy donors with anti-CD3 for 24 hours and sorted CD4+ cells from the stimulated PBMC for RNA-seq analysis. The readings of PTPN22 level obtained from RNA-seq ranged from 4.88 to 38.37 and were comparable between female and male donors (Figure 4A). The age ranged from 29-79 years and there was no correlation between the level of PTPN22 and age either (Figure 4B).

We then arbitrarily categorized all samples into PTPN22hi (>20, N=8) and PTPN22lo (<20, N=14) groups (Figure 4B). We then used Qlucore to identify genes that are differentially expressed between these two groups according to the criteria: q=0.06, fold change <0.25 or >4, and default p value <0.0010271. There were 1035 genes met the criteria (Figure 4C). While PFKFB3, ATM and IL-17F did not come out from the unbiased two-group analysis, the level of PTPN22 positively correlated with the levels of PFKFB3 and ATM, but negatively correlated with that of IL-17F (Figure 4D). This finding is consistent with our published data showing that PTPN22 regulates the expression of PFKFB3, ATM and IL-17F in PBMC [11]. The list of genes was then analyzed with INGENUITY program to identify molecular pathways that were over-represented among the differentially regulated genes and their potential upstream regulators. The top canonical pathways thus identified included EIF2 signaling, telomerase, inosine-5-phosphate biosynthesis, and purine nucleotide de novo biosynthesis (Figure 4E). EIF2 pathway includes many ribosomal proteins that are involved in protein synthesis, whereas inosine-5-phosphate synthesis precedes purine synthesis. Interestingly, the levels of most of the genes listed under EIF2, inosine-5-phosphate biosynthesis and purine synthesis were lower in PTPN22hi group (Figure 4F and 4G), suggesting that PTPN22 negatively regulates protein and DNA synthesis.

The levels of genes listed under the telomerase pathway, including TEP1, TPP1, RAP2B, HDAC5, and ELK3, were in general higher in PTPN22hi group (Supplemental Figure 3). However, only TEP1 and TPP1 are unique to the telomerase pathway; the rest are shared by many other signaling pathways. In addition, upon further reviewing the raw RNA-seq data, we found that TPP1 was indeed tripeptidyl peptidase 1 but not ACD (ACD shelterin complex subunit and telomerase recruitment factor, a.k.a. PIP1 or TPP1). Therefore, the validity of the telomerase pathway is still unclear.

INGENUITY also predicted several upstream regulators of the 1035 genes. The top 4 regulators are CD28, HAVCR1, NR3C1, and Myc (Figure 4H). CD28 is a critical co-activator of T cells. Blocking CD28 signaling with CTLA-4Ig (abatacept) is therapeutic in rheumatoid arthritis, whereas strengthening the CD28 pathway with anti-CTLA4, such as Ipilimumab, is known to cause various autoimmune diseases, including inflammatory arthritis. We therefore postulated that CD28 signals negatively regulates the expression of PTPN22. Interestingly, the level of CD28 itself was actually comparable between PTPN22hi and PTPN22lo groups and did not correlate with the level of PTPN22 either (Supplemental Figure 4). To further examine the role of CD28 in regulating the expression of PTPN22, we stimulated PBMC from healthy donors with anti-CD3 in the presence of anti-CD28. Indeed, anti-CD28 dose-dependently suppressed the expression of PTPN22 (Figure 5A). Anti-CD28 also dose dependently inhibited the expression of ATM, but reciprocally enhanced the expression of IL-17F. Surprisingly, anti-CD28 did not suppress the expression of PFKFB3 but actually slightly increased PFKFB3 level in some donors. When the stimulated PBMC were separated into Th and non-Th cells, we found that anti-CD28 inhibited the expression of PTPN22 and ATM in both Th and non-Th cells (Figure 5B). We subsequently examined whether there was any synergy between CD28 signals and TNFα/TNFαR pathway in regulating the PTPN22 gene signature. Somewhat unexpectedly, we found no substantial synergistic or additive effect between anti-CD28 and exogenous TNFα on the PTPN22 gene signature (Figure 5C).

One possible explanation for the effect of anti-CD28 is that anti-CD28 enhances the expression of endogenous TNFα, which subsequently induces the PTPN22 gene signature. This scenario is unlikely because anti-CD28 failed to suppress the expression of PFKFB3 (Figure 5A). In addition, exogenous TNFα in the absence of anti-CD28 did not affect the expression of PTPN22 in Th cells (Figure 3B). Furthermore, the effect of anti-CD28 on the expression of PTPN22, ATM, and IL-17F was more prominent than that of exogenous TNFα (Figure 5C). Thus, the effect of anti-CD28 is not mediated solely by excessive TNFα induced by anti-CD28.

### 3.6 Counteracting anti-CD28 with TNFα inhibitors

However, it is still possible that the TNFα/TNFαR pathway is essential for anti-CD28 to induce the PTPN22 gene signature, particularly the downregulation of PTPN22 in Th cells, and that endogenous TNFα induced by anti-CD28 is sufficient to provide the assist to anti-CD28. This scenario could explain the lack of synergy between anti-CD28 and exogenous TNFα. Indeed, anti-CD28 further enhanced the expression of TNFα by anti-CD3 stimulated PBMC (Figure 6A). To further test this hypothesis, we stimulated PBMC of healthy donors with anti-CD3 and anti-CD28 in the presence or absence of etanercept or adalimumab. In agreement with our hypothesis, both etanercept and adalimumab neutralized the inhibitory effect of anti-CD28 on the expression of PTPN22 in PBMC (Figure 6B). In addition, when we separated stimulated PBMC into Th and non-Th cells, we found that etanercept and adalimumab also normalized the expression of PTPN22 in Th cells and non-Th cells (Figure 6C). By contrast, we found that tocilizumab, which similar to adalimumab and etanercept also contains the Fc portion of human IgG1 but targets IL-6R, was unable to counteract the effect of anti-CD28 (Figure 6D).

## 4. Discussion

We have demonstrated that a PTPN22-regulated gene signature, which is prevalent in healthy at-risk individuals, is also detected in patients with active RA. This gene signature is reversed after effective treatment or with several targeted therapies in vitro. These findings are consistent with the published data showing a reverse correlation between the level of PFKFB3 and the Disease Activity Score in 28 joints (DAS28) [16]. In addition, ATM is in a gene module of CD4+ T cells that is downregulated in active RA but reversed after abatacept treatment [17]. Furthermore, transcripts of the IL-17 pathway are downregulated in whole blood from responders, but not non-responders, by anti-GMCSFRα [18].

Several groups have tried using synovial tissue pathology and/or “omics” approaches to predict patient’s response to targeted therapies [19–26]. However, these published approaches are often costly, not clinically practical, and not drug-specific. While the data shown in Figure 2 must be validated in larger longitudinal RA cohorts, the in vitro reversal of the PTPN22 gene signature has the potential of becoming a clinically useful and cost-effective assay that can assist rheumatologists in selecting etanercept, adalimumab, or tocilizumab when patients fail methotrexate, bringing us one step closer to personalized treatment of RA.

One intriguing issue is whether the assay can discern response to etanercept versus adalimumab. In the subjects studied in Figure 2A–2D, PBMC were tested with both etanercept and adalimumab. While etanercept and adalimumab yielded concordant results in 9 of the 14 subjects (using 1.4-fold increase in PTPN22 level as the cut-off), discordant results were obtained in the remaining 5 subjects (Table 1). We do not have data on whether etanercept non-responders would have responded to adalimumab or vice versa. Therefore, we were not able to determine if the discordant result correlates with discordant response to these two TNFis. Additional studies with larger sample sizes and including patients how are non-responders are needed to determine whether the in vitro assay is drug-specific and whether it can be used to predict response to other targeted targeted therapies, such as tofacitinib and abatacept.

Our data indicate that exogenous TNFα and CD28 signals partly recapitulate the PTPN22 gene signature through distinct mechanisms in PBMC from healthy individuals. While the details of the mechanisms are still unclear, our data are consistent with reports showing that genetic variations at CD28 are associated with RA [27, 28]. Our data are also very intriguing in light of recent reports of autoimmune diseases, including inflammatory arthritis, induced by immune check-point therapy using anti-CTLA4 in cancer patients. Anti-CTLA4 is expected to enhance CD28 signals and very likely inhibits the expression of PTPN22, thereby leading to the PTPN22 gene signature. However, it remains to be determined if the PTPN22 gene signature observed in active RA is indeed caused by aberrant TNFα and CD28 signals. Unfortunately, we were not able to address this question in this study given the limited supply of PBMC from patients with active RA. In addition, it will be very interesting to examine the anti-CD3-mediated induction of PTPN22 in PBMC obtained from patients received anti-CTLA4.

Neither TNFα nor CD28 signaling fully recapitulate the PTPN22 gene signature, suggesting additional “hits” are required to induce the full PTPN22 gene signature. It is highly likely that the PTPN22 gene signature in each individual patient is caused by a unique combination of those hits. This scenario satisfactorily explains the differential response to targeted therapies in RA patients. The 1000 genes whose expression correlate with that of PTPN22 will provide us with a roadmap for identifying the additional hits triggering the PTPN22 gene signature and additional downstream target genes of PTPN22.

The observation that tocilizumab can reverse the PTPN22 gene signature in responders strongly suggest that IL-6/IL-6R pathway is one of the triggers of the PTPN22 gene signature. However, it is surprising to see that exogenous IL-6 had little impact on the expression of PTPN22. One possible explanation for this unexpected result is trans-signaling triggered by IL-6/soluble IL-6R complex [29]. IL-6 activates STAT3 through membrane IL-6R (mIL-6R) and gp130. While gp130 is a ubiquitous protein, the expression of mIL-6R is restricted to a few subsets of hematopoietic cells. Exogenous IL-6 is expected to act on only the mIL-6R-expressing cells. It has been demonstrated that IL-6 can form a complex with soluble IL-6R (sIL-6R). The IL-6/sIL-6R can then bind to membrane gp130 to activate STAT3 in all cells. Tocilizumab has been shown to disrupt the formation of the IL-6/sIL-6R complex. It is possible that the PTPN22 gene signature is caused by the IL-6/sIL-6R trans-signaling in some RA patients and cannot be recapitulated with exogenous IL-6. This scenario remains to be examined.

It is also somewhat surprising to see that etanercept and adalimumab can counteract the effect of anti-CD28 in Th cells. While one scenario is that endogenous TNFα, which is neutralized by the TNFis, is essential for the effect of anti-CD28, an alternative explanation is reverse signaling mediated by membrane TNFα (mTNFα). Many cells, including T cells, express mTNFα in addition to soluble TNFα. It has been demonstrated that mTNFα, once engaged by TNFα receptor or anti-TNFα, is capable of transmitting signals back into mTNFα-bearing cells [30]. Thus, it is possible that etanercept and adalimumab counteract the effect of anti-CD28 through reverse signaling. This scenario is now under investigation.

## 5. Conclusions

The PTPN22 gene signature is also present in patients with active rheumatoid arthritis and can be reversed after effective treatments. This gene signature can be partially recapitulated by TNFα or CD28 signaling pathways and can also be reversed in vitro with adalimumab, etanercept, or tocilizumab. This in vitro reversal potentially can be used to predict clinical response to the targeted therapies.

## Author Contributions

HHC, CHH, AAS, BT and ICH designed the experiments. HHC, CHH, AAS, and BT carried out the experiments. HHC, CHH, AAS, BT, DAR, YCL and ICH analyzed the data. YCL recruited subjects with active RA and characterized their clinical response to targeted therapies. JAS and EWK identified and recruited healthy donors. DAR collected PBMC and identified suitable donors for this study. HHC, CHH, AAS, BT, YL, DAR, and ICH wrote the manuscript. All authors edited and revised the final manuscript.

## Acknowledgement

We thank Michael Gurish for technical help and Adam Chicoine and the Human Immunology Center Flow Cytometry core for assistance with cell sorting. We also thank Alyssa Wohlfahrt for her assistance in the recruitment of subjects and general oversight of the CPIRA study. We thank the participants and staff of the Partners Biobank for their contributions.

---

**Supplemental Figure 1.**
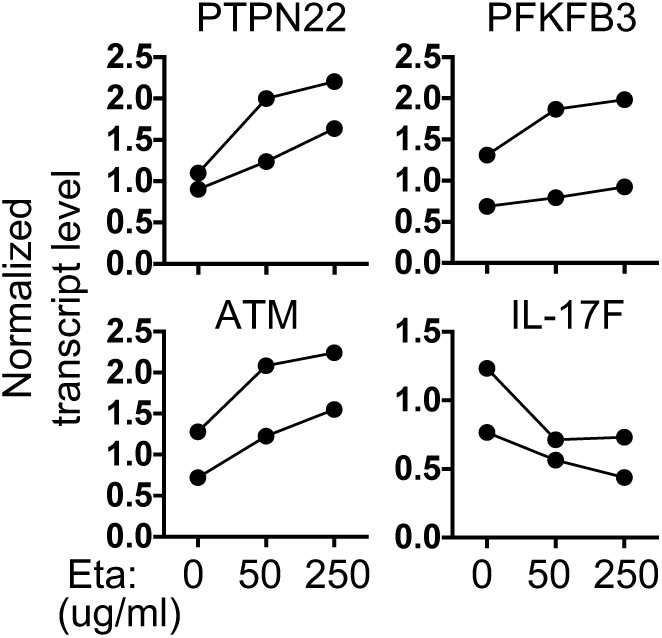
Reversal of the PTPN22 gene signature in vitro by etanercept. PBMC from two patients with active RA were stimulated with anti-CD3 in the presence of etanercept (Eta) at indicated concentrations for 24 horus. The transcript levels of the indicated genes in stimulated cells were measured with qPCR. Anti-CD3 only groups were used as the control groups for normalization of qPCR data. Data points from the same donors were connected with lines.

**Supplemental Figure 2.**
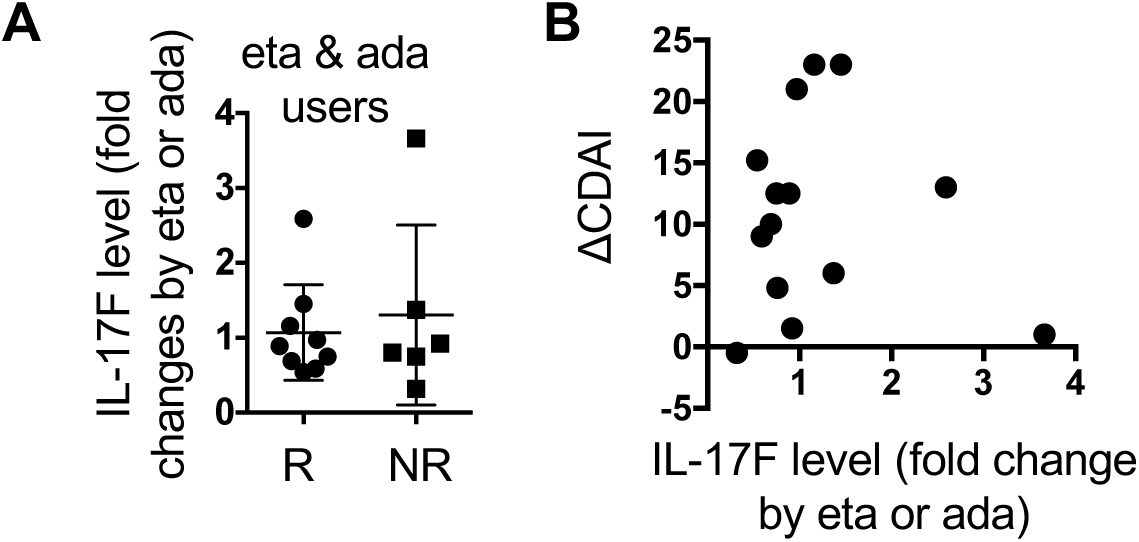
Changes in the expression of IL-17F by etanercept or adalimumab do not correlate with clinical response. The transcript levels of IL-17F from responders and non-responders described in Figure 2 were measured with qPCR (**A**), and plotted against ΔCDAI (**B**).

**Supplemental Figure 3.**
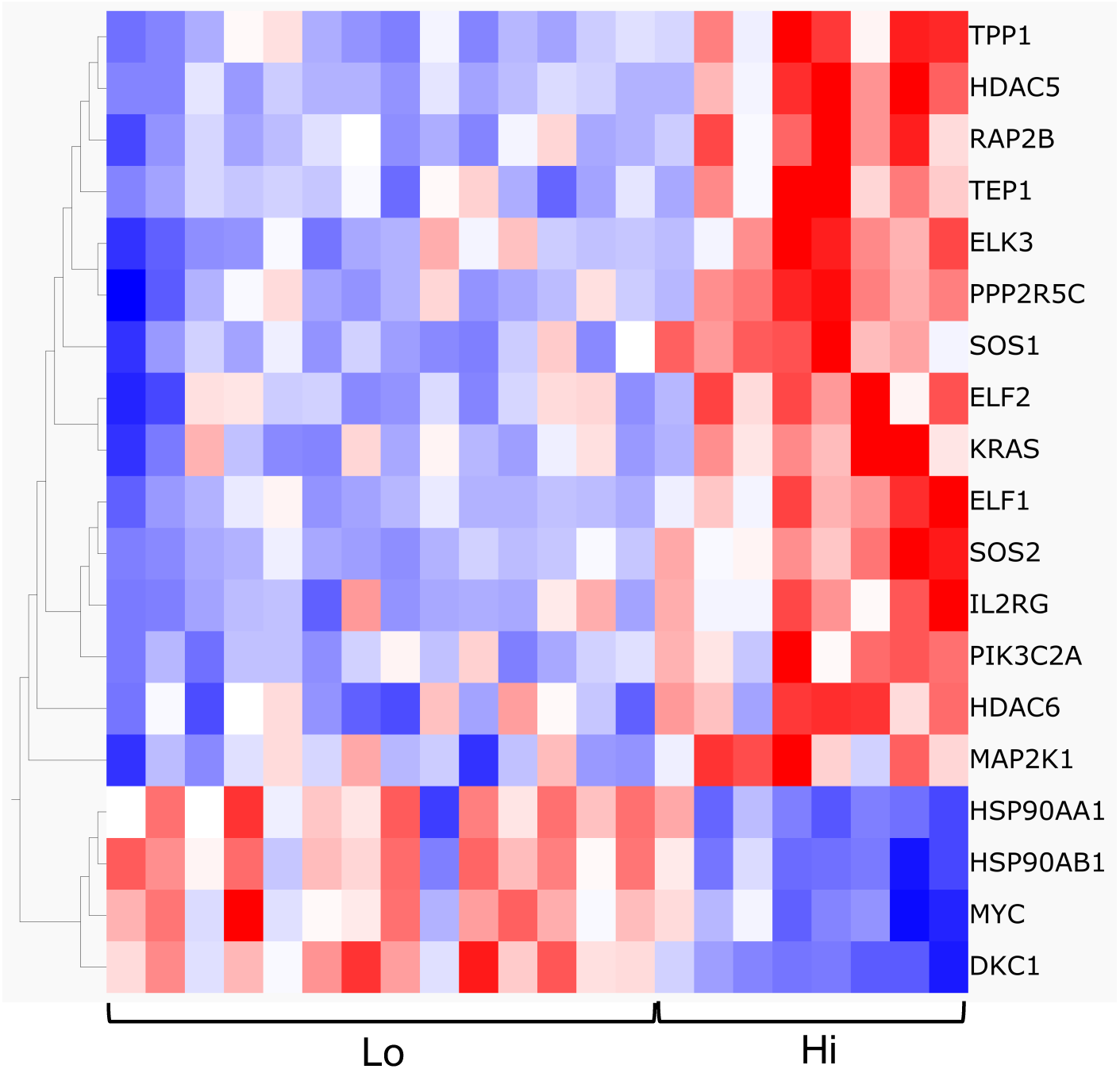
Association between the telomerase pathway and the level of PTPN22. The heat map of the genes listed under the telomerase pathway in PTPN22hi and PTPN22lo group is shown.

**Supplemental Figure 4.**
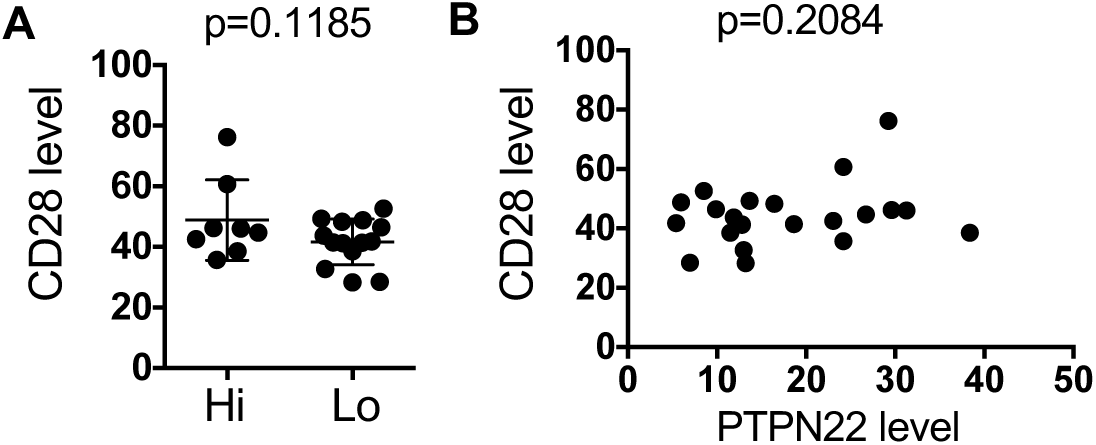
The expression of CD28 does not correlate with that of PTPN22. The levels of CD28 in PTPN22hi and PTPN22lo groups are shown in **A**, or plotted against the levels of PTPN22 in **B**.

**Supplemental Table 1:**
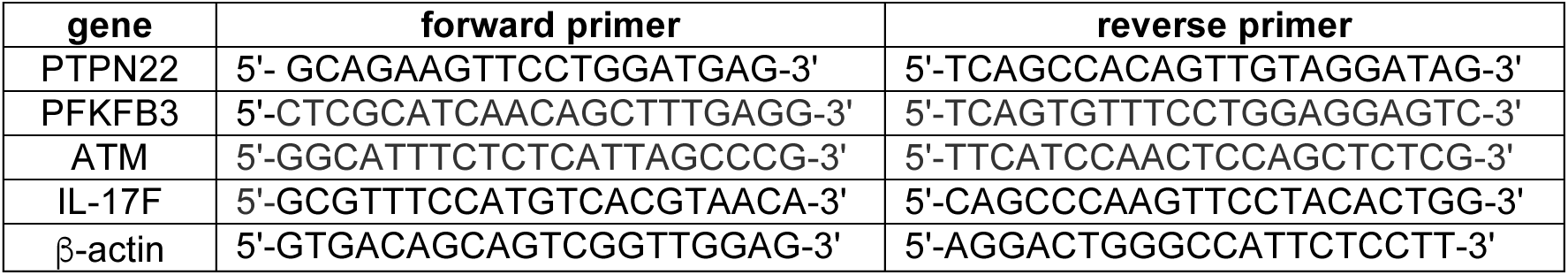
The sequences of the primers used in qPCR

